# Extracting information from RNA SHAPE data: Kalman filtering approach

**DOI:** 10.1101/408872

**Authors:** Sana Vaziri, Patrice Koehl, Sharon Aviran

**Affiliations:** Department of Computer Science, University of California Davis, Davis, California, United States of America; Genome Center, University of California Davis, Davis, California, United States of America; Department of Biomedical Engineering, University of California Davis, Davis, California, United States of America

## Abstract

RNA SHAPE experiments have become important and successful sources of information for RNA structure prediction. In such experiments, chemical reagents are used to probe RNA backbone flexibility at the nucleotide level, which in turn provides information on base pairing and therefore secondary structure. Little is known, however, about the statistics of such SHAPE data. In this work, we explore different representations of noise in SHAPE data and propose a statistically sound framework for extracting reliable reactivity information from multiple SHAPE replicates. Our analyses of RNA SHAPE experiments underscore that a normal noise model is not adequate to represent their data. We propose instead a log-normal representation of noise and discuss its relevance. Under this assumption, we observe that processing simulated SHAPE data by directly averaging different replicates leads to bias. Such bias can be reduced by analyzing the data following a log transformation, either by log-averaging or Kalman filtering. Application of Kalman filtering has the additional advantage that a prior on the nucleotide reactivities can be introduced. We show that the performance of Kalman filtering is then directly dependent on the quality of that prior. We conclude the paper with guidelines on signal processing of RNA SHAPE data.

## Introduction

Beyond its role in protein synthesis and the transfer of genetic information, RNA exists as a dynamic cellular component at the core of gene regulation [1]. From microRNAs involved in regulating gene expression [2] and long noncoding RNAs similarly regulating gene expression [3] to ribozymes acting as chemical catalysts [4], RNA plays a central role in a multitude of cellular activities. The diverse repertoire of biological functions that RNAs adopt is deeply rooted in their abilities to form complex three-dimensional structures [1]. This interplay between structure and function underscores the need for robust structural analysis as a prerequisite to a full understanding of the physiological role of RNA [5]. Despite its importance, determining the complex 3D structures of RNA remains a challenging problem, particularly for longer RNAs [6, 7].

Considering the hierarchical nature of RNA folding [8], much of the efforts in structure determination have been devoted to its two-dimensional base-pairing pattern, also known as its secondary structure. This secondary structure is generally considered to be more stable than and independent of the final 3D conformation [8]. Though experimental methods such as nuclear magnetic resonance (NMR) [9] and crystallography [10] can be used to accurately resolve 3D RNA structures, they are time-consuming, expensive, and often preclude the analysis of long or flexible molecules [11]. Comparative sequence analysis, the process of inferring base-pairing from co-variations observed in the alignment of homologous sequences, is a robust method for defining the secondary structure of RNA [11, 12]. However, this approach has narrow applicability as it relies on the availability of an alignment with a large and diverse set of homologs [13, 14]. An approach that circumvents the need for homologs is *de novo* RNA secondary structure prediction. Many of these sequence-based methods employ a dynamic programming algorithm with a thermodynamics-based scoring function to predict an optimal secondary structure [15, 16]. The resulting computationally predicted secondary structures exhibit variable accuracies [17]. As structure prediction relying on sequence alone poses a difficult problem, the addition of auxiliary experimental data is one way to improve these computational structure predictions [18–20]. The data most commonly included in these prediction algorithms are derived from structure probing experiments [21, 22]. However, little is known about the statistics of these data. One goal of this study is to develop a statistical model for the uncertainty in probing data so that robust information can be extracted.

Structure probing (SP) refers to a class of experiments designed to link chemical reactivity to molecular geometry. In SP experiments, a chemical reagent selectively modifies nucleotides based on their accessibility. In the case of hydroxyl radical experiments, the accessibility is akin to the solvent accessibility [23–25]. Alternatively, in SHAPE (Selective 2’-Hydroxyl Acylation analyzed by Primer Extension) experiments [26], the chemical reagent probes the backbone flexibility of each nucleotide. This flexibility correlates with the pairing state of the nucleotide: higher reactivities are generally observed for unpaired nucleotides. Thus, by extension, the chemical reactivity obtained in such experiments is a probe of the RNA’s secondary structure. In practice, a SHAPE experiment is run as follows: The RNA sample is first modified with the chemical probe. Following this, reverse transcription is applied to detect the resulting chemical modifications along the RNA sequence. Those modifications either cause termination of transcription or introduce a mutation to the transcribed cDNA. Modified locations can then be detected through cDNA fragment sequencing. By comparing to data coming from an untreated control sample, the detection of the modifications is then a direct measure of the reactivity for each nucleotide. The resulting sequence of reactivities is referred to as the reactivity profile or simply the *profile* of the RNA. Recent advances in sequencing have ushered in a new era of affordable and massively parallel SP experiments [20] and applications of resulting data are not limited to structure prediction. In fact, among other uses, SP has been used to direct sequence alignment as well as to strengthen evolutionary signals when searching for conserved RNA structures between organisms [27].

As with any measured value, SP reactivities are corrupted by noise. The standard approach taken by experimentalists to reduce the impact of that noise is to repeat the experiment multiple times under the same condition and combine the results using basic averaging. For SP data, we use the term *replicates* to refer to the multiple reactivity profiles and *measurements* to refer to the set of reactivities for a particular nucleotide coming from these replicates. Basic replicate averaging is performed by taking a per-nucleotide average across measurements. This sequence of values forms the average profile. While straightforward, this implies that the noise is additive and has zero-mean. These criteria have not been established for SP experiments. Indeed, the noise observed in SP data has not been explicitly studied and currently no models exist to characterize the observed differences between replicates. In this manuscript, we propose a model for the noise associated with SHAPE data and develop a pragmatic approach to signal denoising. To this end, we borrow from the comprehensive literature available on denoising in signal and image processing (see for example [28] and [29]). We first note that previous analysis of SHAPE data has revealed the log-normality of reactivities [30]. This observation naturally led us to study the replicate noise after applying a logarithmic transformation. Log transformations of data are simple and easily reversible operations that are often applied in the case of skewed data to mitigate the effects of volatile measurements [31]. Apart from their extensive use in image processing, they have also been widely studied in the context of biological data analysis, such as in microarray data analysis where they can act as a variance stabilizer [32]. It is worth mentioning that in dynamic programming based secondary structure prediction methods, such as [18], SHAPE data are integrated into the prediction algorithm via a logarithmic relationship between the reactivities and a pseudo-energy term. This operation implicitly decreases the impact of nucleotides with high reactivity [33]. In this work, we propose an additive Gaussian noise model for log transformed SHAPE data. This transformation allows us to study signal processing techniques that leverage the log-normality of the SHAPE distribution as prior information. In particular, we apply the Kalman filter [34, 35], an algorithm commonly used in signal processing and control theory, to SHAPE data. This filter works by optimally fusing two sources of information: prior knowledge on nucleotide reactivity and the noisy measurements. It has previously been applied to protein structure determination from NMR data [36, 37]. For our purposes, we use the log-normal distribution of SHAPE reactivities as the required prior with the goal of optimally extracting true reactivity information from the noisy measurements.

In this work, we explore the following questions. First, how much of an advantage over averaging does a sophisticated denoising strategy, such as Kalman filtering, offer when extracting a reactivity signal from noisy replicates? Second, how many replicates are required for robust signal extraction? Given that the majority of published SP data consists of between one and three replicates, these questions are critical to experimental design. We address these questions under the assumptions of our proposed noise model. The paper is organized as follows: In the Background section, we provide an overview of SHAPE experiments followed by a discussion on the factors contributing to noise in these experiments. We then discuss important characteristics of SHAPE data and give a brief overview of signal filtering. In the section that follows, we revisit the statistical models used in replicate processing and propose a noise model based on the log transformation. We then provide a description of how Kalman filtering can be applied as a denoising strategy in the context of replicate processing. In the Results section, we compare the approaches of averaging and Kalman filtering using replicates simulated under the proposed statistical model. Finally, we conclude with a discussion on the statistical models and signal processing methods described and future directions.

## Background

### Overview of SHAPE experiments and reactivity reconstruction

In a typical SHAPE experiment, a sample of an RNA is treated with a chemical reagent that selectively forms adducts on nucleotides along flexible regions of the molecule. After treatment, reverse transcription is applied to detect locations of adduct formation. The adducts interfere with this transcription, either by causing termination or, in the case of SHAPE-MaP experiments [38], by introducing a mutation in the nascent cDNA strand. Lengths of the cDNA fragments, or equivalently, mutation sites, correspond to their locations along the RNA. The number of modifications per nucleotide are then converted into a modification rate. Reverse transcription is simultaneously applied to an untreated sample of the RNA. One way to determine a reactivity value per nucleotide is to compute the difference between the modification rates per-site on the reagent-treated and control samples [39, 40]. The reactivity resulting from this background-subtraction is a measure of the nucleotide’s sensitivity towards the reagent and correlates with the local backbone flexibility [41]. As structurally constrained regions of an RNA correspond to base-paired nucleotides, nucleotides exhibiting low reactivities are likely paired while highly reactive nucleotides are indicative of unpaired regions of the RNA [19].

Prior to use in further analysis, reactivity profiles are normalized so that values across a transcript span the typical range of values between 0 and 2. This is done to ensure uniformity between replicates as well as across different transcripts [33]. One commonly applied model-free normalization technique works as follows: First, a percentage of the data corresponding to the highest reactivity values are considered outliers and are removed from the analysis. According to [19], for RNAs shorter than 100 nucleotides, no more than 5% of the data should be removed and for longer RNAs, no more than 10% of the data. From the remaining nucleotides, another band of highly reactive nucleotides (usually around the top 8-10%) are averaged in order to calculate a normalization factor [19, 33, 42]. The entire profile, including the previously excluded outliers, is then normalized by this factor. On the normalized scale, reactive nucleotides are roughly defined as those with reactivities higher than 0.7 and unreactive nucleotides are those with reactivities below 0.3 [18].

Currently, there is no standardized practice for normalization [40] and, even after normalization, it is not uncommon to observe values significantly higher than 2. Additionally, while the standard values of reactivities are positive, negative-valued reactivities are often observed in the data. These values occur when there is a stronger readout in the control sample compared to the reagent-treated sample and the background-subtraction process does not completely account for sequence-specific noise. In practice, negative values are simply set to 0 [33].

### Factors contributing to variation in SHAPE experiments

There are a number of influencing factors when it comes to uncertainty in SHAPE reactivity values. Discrepancies observed between replicates can be classified as stemming from two main sources [40].

The first source of noise can be classified as *technical variation* and includes anything from the stochasticities introduced by the sequencing platform to the multiple steps in the cDNA library preparation. Technical considerations also include variations that are a product of the dynamic nature of RNA: RNAs in a sample can fold into and transition between various structures. These changes are sensitive to numerous parameters involved in the probing experiment, including solvent conditions, temperature, and protein interactions [43]. As SHAPE reactivities represent an aggregate measure on all RNA copies co-existing in a sample [44], parametric fluctuations ultimately manifest as observable differences between replicates. RNA thermometers, which shift from a highly structured state to an unfolded state with increasing temperature, are one clear example that demonstrate this effect [45]. The relative concentrations of the two states ultimately cause temperature-dependent variations in the measured reactivities.

Along with technical factors, inter-replicate divergences can also be caused by biological factors in the underlying sample. Such effects are referred to as *biological variation*. One example is the degree of structural diversity in the sample being probed. It is known that the same RNA sequence can fold into many different structures that co-exist with varying abundances in a sample. Riboswitches, for example, are RNA elements whose functionality hinges on their ability to alternate between two conformations to regulate gene expression [46]. This switching between folds cannot be instantaneous without violating physical laws: the change in structure must be gradual and thus gives way to the existence of intermediate structures between folding pathways. As a SHAPE reactivity reflects the combined reactivity of all RNA copies co-existing in the sample, the degree of structural diversity in the sample ultimately affects the differences between replicate measurements.

The discrepancies between replicates reflect a composite effect of both the technical and biological variation. We refer to this combination as the *measurement noise*, which we aim to model.

### Characteristics of SHAPE data

As SHAPE profiles include measurement noise, a term we use to span the effects of multiple facets of experimental uncertainty, any analysis of these data must include a denoising step. The traditional approach is to compute the average across replicates. This method is sensible under an implicit assumption that the true reactivity value of a nucleotide is corrupted by additive noise that follows a zero-mean distribution. Most often, this distribution is assumed to be Gaussian. However, the number of processing steps involved in the quantification of the SHAPE profile, namely, computing the chemical modification rates, the background-subtraction, and the normalization processes, raise doubts about this assumption. We diverge from the traditional approach and propose a log transformation based noise model that renders the data amenable to well-established signal processing techniques. The foundation of our noise model, which will be introduced in the following section, was further prompted by the following fundamental observation on SHAPE data: the empirical distribution of SHAPE reactivities is highly skewed. This distribution is in fact near-Gaussian after applying a log transformation [30]. We adopt this log-normality as an assumption for the remainder of our work.

Before proceeding, we note that some caution is required when defining a noise model for SHAPE data for the following reasons. First, the normalization of SHAPE reactivities does not preclude negative values and such reactivities are incompatible with the log-normal model. While negative values are not rare, they are assumed to occur when the control sample can not be used to adequately describe the true noise component of the reagent-treated sample. In this case, subtraction of the control modification rate from the reagent-treated rate does not suffice as a correction method. As mentioned, negatives are commonly set to 0 in the final SHAPE profile. This practice skews the distribution for unreactive nucleotides and strongly implies an asymmetric distribution of measurement noise.

Second, it has been documented that the highest SHAPE values exhibit the most variability between measurements [33]. This is particularly noteworthy as normalized profiles often exhibit highly reactive nucleotides. The relationship between the average reactivity value for a nucleotide across measurements and the standard deviation between measurements reveals the heteroskedastic nature of SHAPE data. Fig 1 illustrates the strong mean-dependence present in the standard deviation values across 5 experimental replicates obtained for the RNA3 segment of the cucumber mosaic virus genome [47]. The log of the measurement standard deviations and log of the measurement means are related linearly and the slope of this relationship nearly 1. Equivalently, the measurement standard deviation is nearly proportional to the measurement mean, which may be indicative of a multiplicative noise term. Thus, the standard statistical model relying on an assumption of an additive noise term may not properly serve SHAPE measurements.

**Fig 1.**
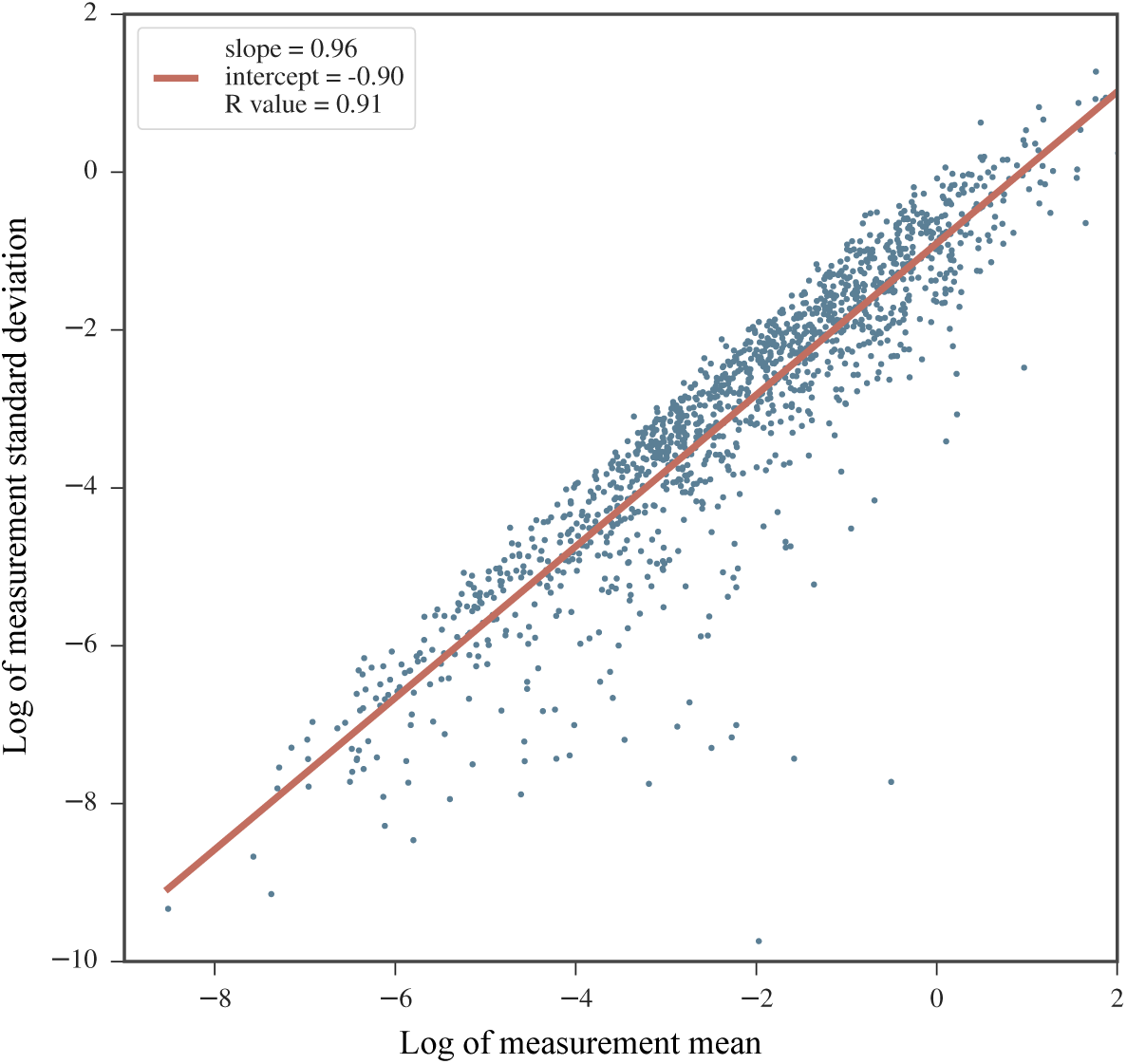
The mean-dependence in the standard deviation of SHAPE measurements. Data from 5 SHAPE replicates obtained on the cucumber mosaic virus RNA3 sequence (experiments performed on data from infected plant cell lysates) [47]. For each nucleotide, the mean value of the 5 measurements were calculated and plotted against their standard deviation on a log-log plot. A linear fit is overlaid in red. Note that negative reactivity values were not included as they are incompatible with the log-log plot.

The two extremes of SHAPE reactivities discussed, namely, those corresponding to unreactive and highly reactive nucleotides, underscore the unique characteristics of SHAPE data. Along with the log-normality of the SHAPE distribution, these characteristics prompted our study of a noise model that relies on a log transformation.

### Signal filtering

The purpose of filtering is to deduce meaningful information from a signal containing unwanted components. Filtering usually relies on the availability of multiple realizations of the signal. The simplest and most common filtering approach utilized in experimental studies is to average the data realizations. Such a filtering relies heavily on the assumption of an inherent randomness in the noise that can be modeled as independent samples of an additive Gaussian distribution. Averaging, however, is not the only form of filtering available from signal processing. In fact, it may not be optimal if the assumption of additive Gaussian noise is invalid. A filter is optimal if it produces the best estimate under a certain prescribed criterion or model [48]. One example of an optimal filter is the Kalman filter (KF) which estimates a parameter in a system affected by additive Gaussian noise. This filter is often utilized in optimal tracking systems and signal processing problems to smooth noisy data or to estimate a parameter from a set of noisy measurements [49]. For the KF, the optimality criterion is defined as minimization of the mean-square error associated with the parameter estimate. At a high-level, the 1 dimensional KF works by iterating between the following two steps:

1. **Predict**: the filter makes a prediction for the current state of the system on which measurements are being made. This prediction is based on a model describing the state dynamics. During the primary predict step, an initial prior on the system state is required to estimate the state sans measurements.
2. **Update**: upon receiving new information in the form of a noisy measurement, the state model is updated. A quantity known as the *Kalman gain* is calculated and is used to optimally combine information from the prior and the newly incorporated measurement. The state model is updated conditioned on the new measurements using the Kalman gain. The updated conditional distribution is then used as a prior distribution in the ensuing predict step.

The Kalman gain is an optimal weighting factor between the previous prediction and the newly observed measurement. Its value depends on the uncertainties of both the prediction and the new measurement. Initially, the prediction is based solely on the input prior. When the measurement is noisy, the model relies more heavily on the prior. Conversely, when the measurements are reliable, the filter puts less weight on the prior. After all measurements have been handled, the final prediction is taken as an estimate of the parameter of interest. This prediction represents an optimal fusion of the prior and the measured values. In classical Kalman filtering applications, the input data is a discrete time series of measurements on a system in which there are two sources of uncertainties: 1. the model dictating the state of the system and its dynamics and 2. the measurements at each time point. For those interested in a derivation of the complete filter and proof of its optimality, we recommend reading [35, 49, 50]. For our purposes, the “state” of the system is a nucleotide’s true reactivity value. The measurements are taken directly on this reactivity and are corrupted by noise. Our aim is to remove the errors in these measurements and recover the true reactivity. A full mathematical characterization of the KF implementation employed in this work is provided in Methods.

## Models for signal extraction in SHAPE data

Below, we introduce notation and discuss two noise models for SHAPE data. We also review the methods used for signal extraction under each model.

### Notation

Consider data coming from *N* repeated SHAPE experiments on an RNA with *M* nucleotides. For each nucleotide *m*, we assume an underlying ground truth reactivity value denoted *s*_*m*_. The sequence of ground truth reactivities making up the true profile is denoted by *S*. The *N* measurements of *s*_*m*_ are denoted 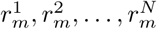. After a log transformation, the measurements are denoted 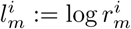. We refer to these values as *log measurements*. Similarly, *l*_*m*_ := log *s*_*m*_ denotes the log of the nucleotide’s ground truth reactivity, or its *log reactivity*. We say the transformed data is in the *log domain* while the original data is in the *data domain*. The sequence of log-transformed ground truth reactivities is denoted *L*. Our goal is to combine the measurement values for each nucleotide in a manner that optimally extracts the true reactivity. This amounts to either recovering *s*_*m*_ from the 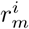 values in the data domain, or, equivalently, *l*_*m*_ from the 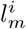 values in the log domain.

### Normal Noise Model

Measurements across replicates for a nucleotide are generally combined into a single reactivity by taking their average. This naive combination is appropriate if the assumed relationship between the *i*^th^ replicate 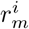 and the ground truth reactivity *s*_*m*_ is governed by the following relationship:

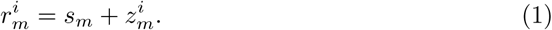

Here, 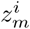 is the measurement noise term, which is assumed to follow a zero-mean Gaussian distribution with standard deviation 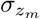 We term this model the *normal noise model*. Under this model, the average reactivity for a nucleotide is

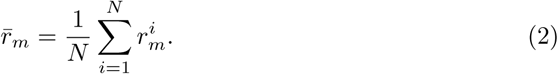

Assuming independence in the 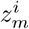 s, this is also the maximum likelihood estimate for *s*_*m*_ [51]. *We refer to the sequence of M* nucleotides averaged in this way as the *average profile* and denote it 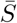. Although it is often not explicit, data processing pipelines that employ an average across measured values are predicated on such a normal noise model. Despite being a straightforward approach to combining replicates, averaging in this way relies on a key assumption of the normal noise model that has yet to be experimentally verified; that is, the assumption of an additive Gaussian distribution of noise in the data domain for probing data.

### Log-Normal Noise Model

We have discussed three noteworthy features of SHAPE data: its log-normal distribution, the skew in measurements introduced by replacing negative-valued reactivities with zeros, and the heteroskedasticity observed in replicates. These features allude to an asymmetric noise distribution. As the empirical SHAPE distribution is Gaussian in the log domain, it is a natural extension to assume that the noise in measurements follows a similar distribution. We were thus motivated to study the data after a log transformation and further modeled the noise as following an additive Gaussian distribution in the log domain. In such a model, the log measurement 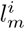 is related to the ground truth *l*_*m*_ according to the following relationship:

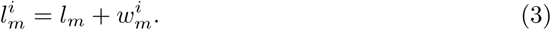

The measurement noise term, 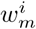, is assumed to follow a zero-mean Gaussian distribution with standard deviation 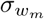. The 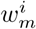 values are assumed to be independent between measurements. We refer to this model as the *log-normal noise model*. As before, the log measurements can be combined by taking their average. To distinguish it from averaging in the data domain, we will refer to this process as *log-averaging*. The log-averaged estimate of *l*_*m*_ is

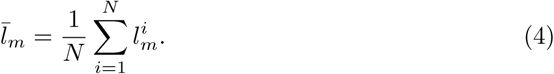

By reverting back to the data domain, we obtain 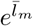 as the final estimate for the reactivity *s*_*m*_. In the log domain, the sequence of log-average reactivities for the *M* nucleotides is denoted 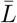. After reverting to the data domain, we refer to the sequence of log-average reactivities as the *log-average profile* and denote it 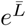. We note that additive noise in the log domain implies multiplicative noise in the data domain, hence

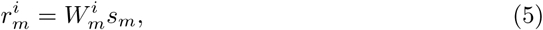

where 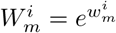.

The central assumptions of the log-normal noise model render the problem of optimally extracting a reactivity value from noisy measurements directly applicable to Kalman filtering. The KF exploits the distribution of the SHAPE data in the log domain as an auxiliary information source and uses it to extract information from noisy measurements. We apply a simplified version of the 1 dimensional KF to a system consisting of a single nucleotide with a ground truth reactivity value that persists between measurements. The measurements of the system state, i.e. the nucleotide’s reactivity, are described by Eq. 3. The filtering process is carried out in the log domain separately for each nucleotide. The KF inputs are summarized below:

1. The log measurements, 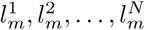, which make up the measurement vector.
2. The uncertainty in the measurements, 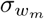. This value is estimated using the sample variance of the 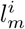 values. It is required by the filter to calculate the Kalman gain.
3. The empirical distribution of log-transformed SHAPE data fit to a Gaussian distribution, *𝒩* (*µ*_0_, *σ*_0_). This is used as the prior in the initial predict step.

The resulting KF reactivity is denoted *k*_*m*_ and is an estimate of the log reactivity, *l*_*m*_. Transforming back to the data domain gives 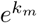 as an estimate of the reactivity, *s*_*m*_. The sequence of filtered reactivities is denoted *K* in the log domain and *e*^*K*^ in the data domain. We refer to *e*^*K*^ as the *Kalman filter profile* or *KF profile*. A detailed description of our KF implementation is provided in Methods. The two log domain processing pipelines, log-averaging and Kalman filtering, are summarized in Fig 2.

**Fig 2.**
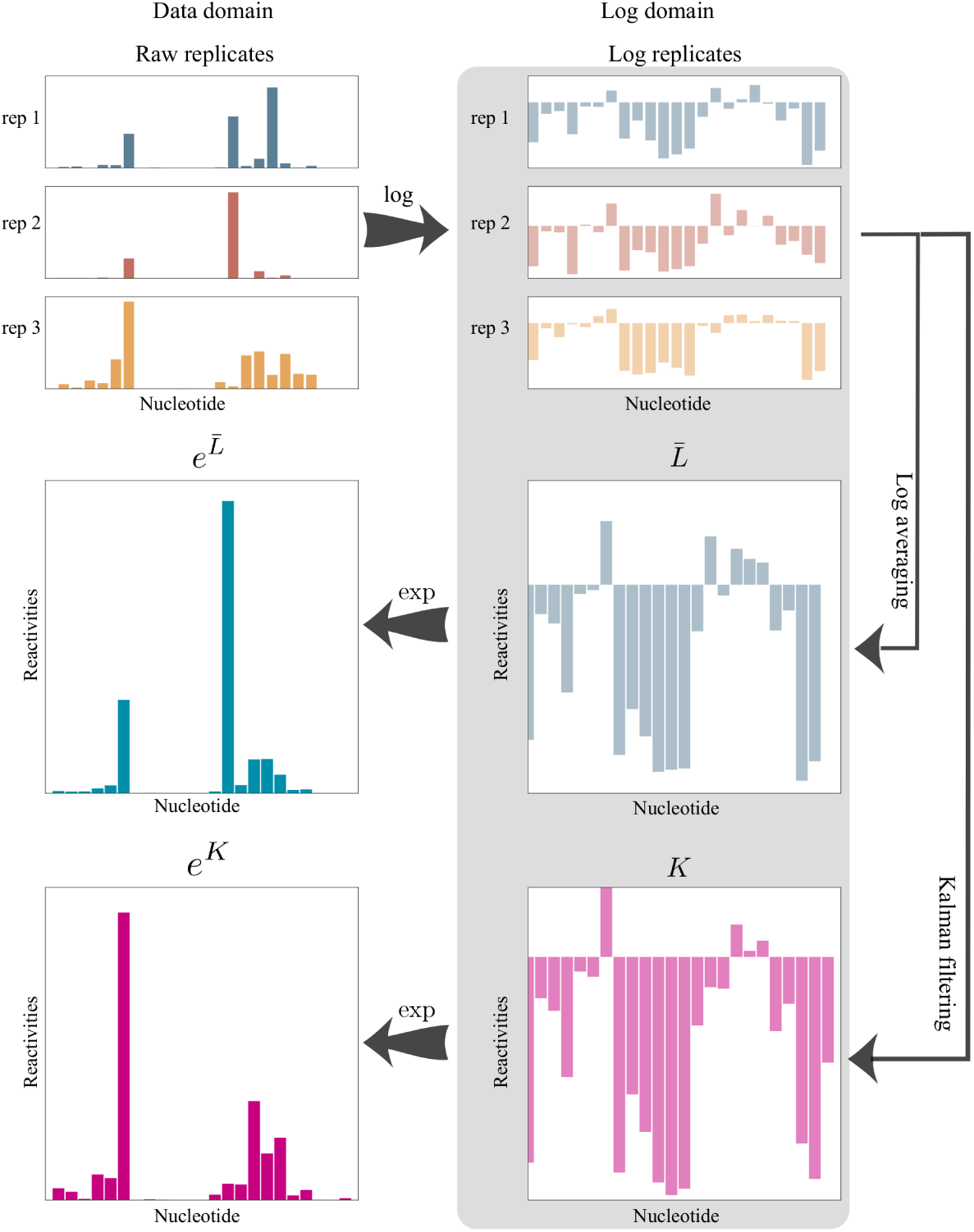
A conceptual representation of measurement combination methods under the log-normal noise model for three SHAPE replicates. Replicates are first transformed into the log domain. The log-average (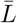) and KF (*K*) profiles are then computed. The resulting profiles are transformed back to the data domain.

## Results

We compared the two statistical filtering approaches for analyzing SHAPE replicates in the log domain introduced above: log-averaging and Kalman filtering. The results presented below are organized as follows. First, we discuss noise levels that are observed in real SHAPE experiments. Then, using simulations, we compare the accuracies of log average profiles to KF profiles by evaluating the ability of each approach to recover the ground truth profile. Finally, we compare data-directed secondary structure prediction results on profiles processed under assumptions of the normal and log-normal noise models.

### Noise levels observed in SHAPE experiments

We studied the noise observed in SHAPE data collected on the 2216 nucleotide RNA3 segment of the cucumber mosaic virus [47]. Included in this analysis were data coming from experiments run on three forms of the RNA: *in vitro* (5 replicates), purified viral RNA extracted from virion particles (3 replicates), and from infected plant cell lysates (3 replicates). Measurements were first transformed to the log domain. We then calculated the sample standard deviations of the log measurements per nucleotide for each of three different forms of the RNA. Thus, standard deviation values were calculated using either 3 or 5 measurements. A histogram of these values and their empirical cumulative density function (CDF) are shown in Fig 3. We used these data to define low, medium, and high noise regimes as follows:

**Fig 3.**
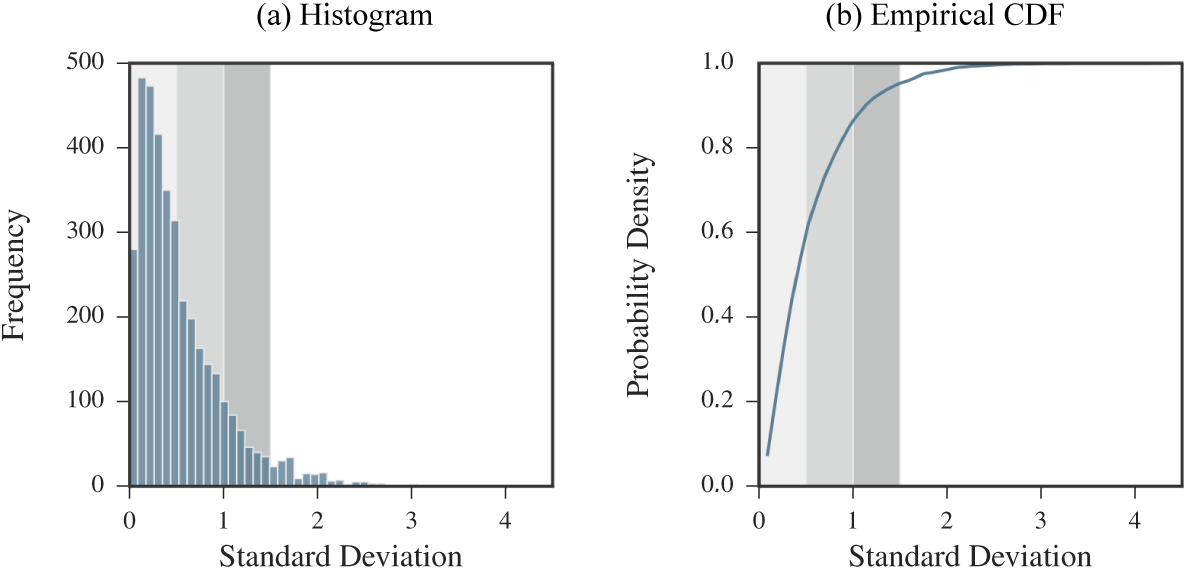
Log domain standard deviation values of measurements coming from real SHAPE data. Standard deviation values were calculated for each nucleotide on log measurements. (a) Histogram of standard deviation values. (b) Empirical CDF of standard deviation values. The shaded regions correspond to our definition of low, medium, and high noise regimes.

All non-positive measurements were removed from the initial set of data. Nucleotides with a single positive measurement were excluded so that a total of 3723 data points were considered.

1. We defined the **low noise regime** by measurements with log domain standard deviation values between 0.5 and 1. This corresponds to about 60% of the data with log domain standard deviation values lying in the 60^th^ percentile.
2. We defined the **medium noise regime** by measurements with log domain standard deviation values between 0.5 and 1. This range was selected to lie between the low and high noise regimes and covers about 26.5% of the data.
3. We defined the **high noise regime** by measurements with log domain standard deviation values between 1 and 1.5. This corresponds to about 10% of the data, with log domain standard deviation values lying between the 86.6^th^ and 95.3^rd^ percentile of the data.

Based on these ranges, we simulated replicates in the log domain with different noise levels by uniformly selecting a standard deviation value within one of the specified ranges (either low, medium, or high). Note that just under 5% of the nucleotides in this analysis exhibit variability in measurements exceeding the high level.

### Kalman filter improves information extraction from noisy replicates

We compared the performances of log-averaging and Kalman filtering for replicates simulated under the log-normal noise model. We first assembled a database of 22 RNAs with published SHAPE profiles and reference secondary structures [18, 30, 52–54]. The database includes ribosomal RNAs, riboswitches, and viruses. RNA lengths vary from 34 to 2094 nucleotides and sum to a total of 11070 nucleotides (see Table 1 of Methods for a complete description). The known SHAPE profiles were treated as ground truth. We simulated 3 replicates for each sequence according to the log-normal noise model. We varied the simulated noise level by increasing the standard deviation of the log measurements from 0 to 5. We then assessed the signal extraction capabilities of log-averaging and Kalman filtering by comparing each resulting processed reactivity to the ground truth. Root mean square (RMS) errors for varying reactivity and noise-levels are shown in Fig 4 (a). In low noise regimes, the two methods performed comparably. However, in higher noise regimes, Kalman filtering recovered better the ground truth reactivity than did log-averaging.

**Fig 4.**
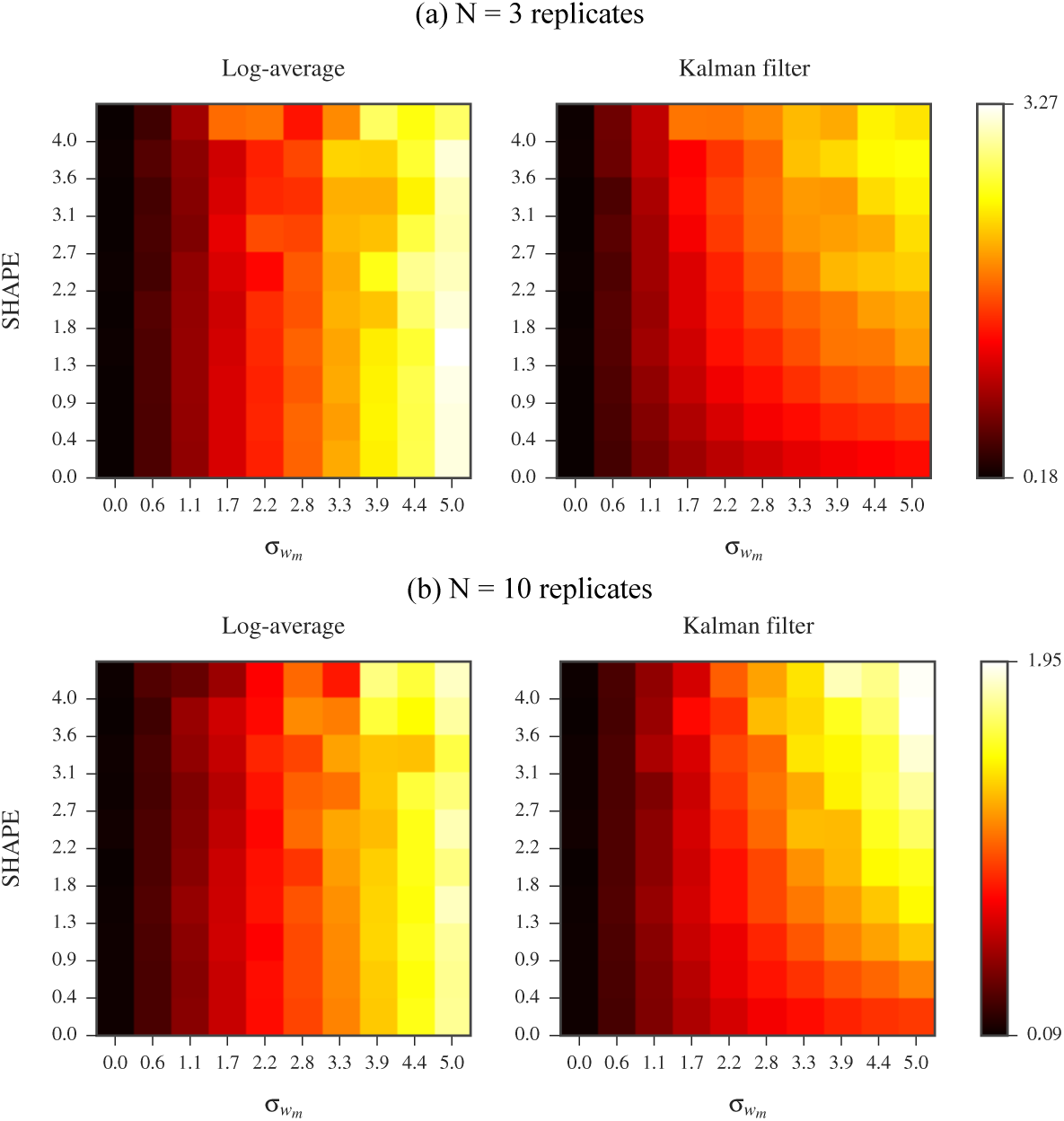
Comparison of log-averaging and Kalman filtering using. **(a)** *N* = 3 **and (b)** *N* = 10 **simulated replicates under log-normal noise model.** The vertical axis represents the data domain ground truth reactivity, *s*_*m*_. The horizontal axis represents the log domain standard deviation of the simulated measurements, 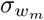 Nucleotides were binned based on *s*_*m*_ and 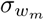 values. Left panel shows RMS errors calculated between ground truth and log-averaged reactivities for all nucleotides in a bin. Right panel shows RMS errors calculated between ground truth and Kalman filtered reactivities for all nucleotides in a bin. Error calculations were carried out in the log domain and ground truth values were the log reactivities. See Methods for RMS calculation details.

We repeated this analysis using 10 simulated replicates for each RNA. The RMS errors for the two processing methods are shown in Fig 4 (b). With this increase in replicates, as expected, both methods exhibited an increase in performance compared to using 3 replicates. Additionally, the simple log-averaging estimate extracted the true reactivity profile as accurately as the more complex Kalman filtering approach, even in the higher noise regime. Hence, Kalman filtering is a more robust method for signal extraction in the case of high noise levels or limited replicates.

### Using more than four replicates marginally improves accuracy

The results presented in the previous section emphasized the impact of replicate count on the relative performances of log-averaging and Kalman filtering. Given 10 replicates, the accuracy of the log-averaging approach mirrors that of Kalman filtering, even in the presence of substantial noise. However, 10 experimental replicates are almost never obtained in practice. To explore how the accuracies of both approaches are affected by replicate count, we repeated our simulations using from 2 to 10 replicates and performed log-averaging and Kalman filtering for each replicate count. We performed this simulation for replicates generated at low, medium, and high noise levels for all RNAs in our database. The RMS errors for both methods are shown in Fig 5 plotted against the number of replicates. These plots reinforces the results presented above: for moderate noise, log-averaging and Kalman filtering perform comparably. Meanwhile, in the high noise regimes, Kalman filtering better recovers the ground truth. This advantage is only present for a small number of replicates, specifically, less than 4. If the number of replicates is increased above this, then the two methods perform comparably even in the presence of high noise. Thus, increasing the number of replicates to be more than 4 does not significantly improve the results of either method. Based on these findings, we recommend obtaining a minimum of 4 replicates.

**Fig 5.**
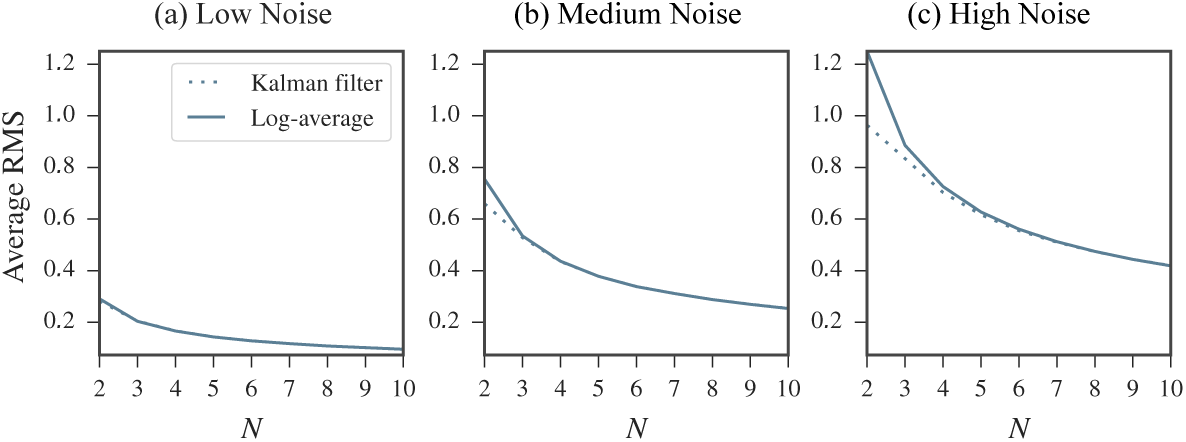
Comparison of the log-average and Kalman filter approaches using *N* = 2 to *N* = 10 replicates simulated at (a) low (b) medium and (c) high noise levels under log-normal noise model. RMS errors were calculated between ground truth and log-averaged reactivities (solid line) and between ground truth and the Kalman filtered reactivities (dotted line) over entire set of nucleotides. Error calculations were carried out in the log domain and the ground truth values were the log reactivities. See Methods for RMS calculation details. In moderate noise regimes, only a negligible difference between the log-averaging and Kalman filtering approaches is observed. However, in the high noise regime, the Kalman filtering approach better recovers the ground truth. This advantage is marginal after the replicate count is increased beyond 4. Note that errors increase with increasing noise levels.

### Refining the Kalman filter prior improves accuracy

The results of the log-averaging approach can be improved either by increasing the number of replicates or by improving the data quality. In contrast, Kalman filtering offers an additional channel for improvement by way of the prior distribution. The prior is used by the filter along with the measurements to extract signal information. Thus, the success of the KF relies on how faithful this model is to the data, in addition to the data quality. With a well-tailored prior, we expect an improvement in Kalman filtering results. Here, we demonstrate this idea with a simple simulation in which we defined an “ideal” prior specialized for each nucleotide. This ideal prior is a Gaussian distribution centered at the ground truth (log reactivity) for that nucleotide and with a small standard deviation. We studied how deviations for this ideal prior affected the KF results by examining the effects of two possible changes. The first was a shift in the prior mean away from the ground truth. This mean offset represents a loss of accuracy in the prior. The second was an increase in the prior standard deviation, representing a loss of precision in the prior. The definitions of the ideal prior and the deviations are described in detail in Methods. We calculated the Kalman filtered reactivity with different mean offset and standard deviation values for 3 replicates simulated under the low, medium, and high noise regimes. The RMS errors calculated over all nucleotides in our database are shown in Fig 6. As this result confirms, the quality of the KF results are related to that of the prior. The KF applied with a prior having high accuracy and precision performs the best.

**Fig 6.**
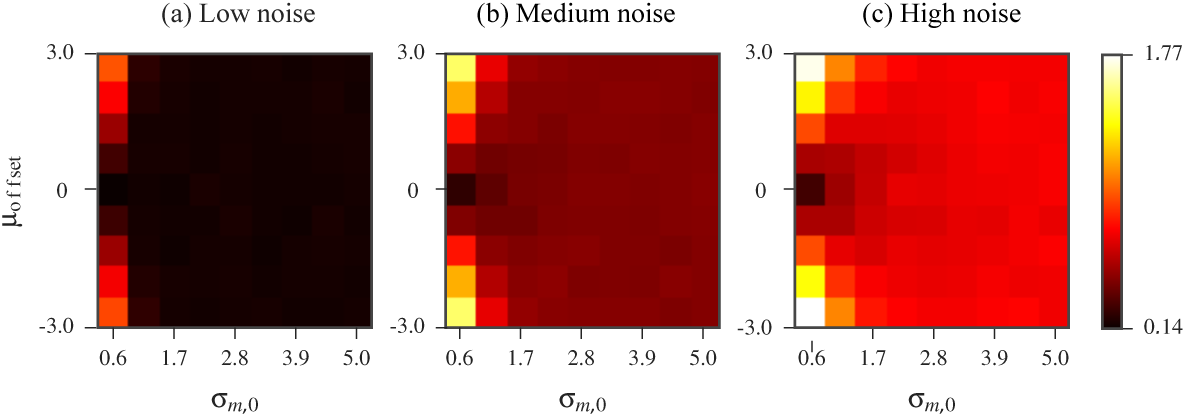
KF results as the prior mean and standard deviation are varied for *N* = 3 replicates simulated at (a) low (b) medium and (c) high noise levels under log-normal noise model. The horizontal axis represents an increase in the prior standard deviation, *σ*_*m,*0_. The vertical axis represents the offset, *µ*_offset_, which was added to the ground truth log reactivity to define the prior mean. The value of each bin is the RMS error calculated over all nucleotides in our database between the ground truth and Kalman filtered reactivities. Error calculations were carried out in the log domain and the ground truth values were the log reactivities. See Methods for RMS calculation details.

Intuitively, applying the KF with a prior that is inaccurate (i.e. having a large mean offset) and precise (i.e. having a small standard deviation) results in the filter placing a high level of confidence in a biased initial prediction. On the other hand, applying the KF with a prior that is inaccurate but also imprecise (i.e. having a large standard deviation) is comparable in performance to the log-averaging approach. This is because the KF places a high level of confidence in the measurements while the prior is largely ignored. To confirm this intuition, we performed the following two experiments:

- The prior used had a mean that was offset from the ideal by a fixed value. We increased its standard deviation and studied the effects on the KF results. RMS errors are shown in Fig 7 plotted against the prior standard deviation.
- The prior used had mean that was fixed at the ideal value. We increased its standard deviation and studied the effects on the KF results. RMS errors are shown in Fig 8 plotted against the prior standard deviation.

**Fig 7.**
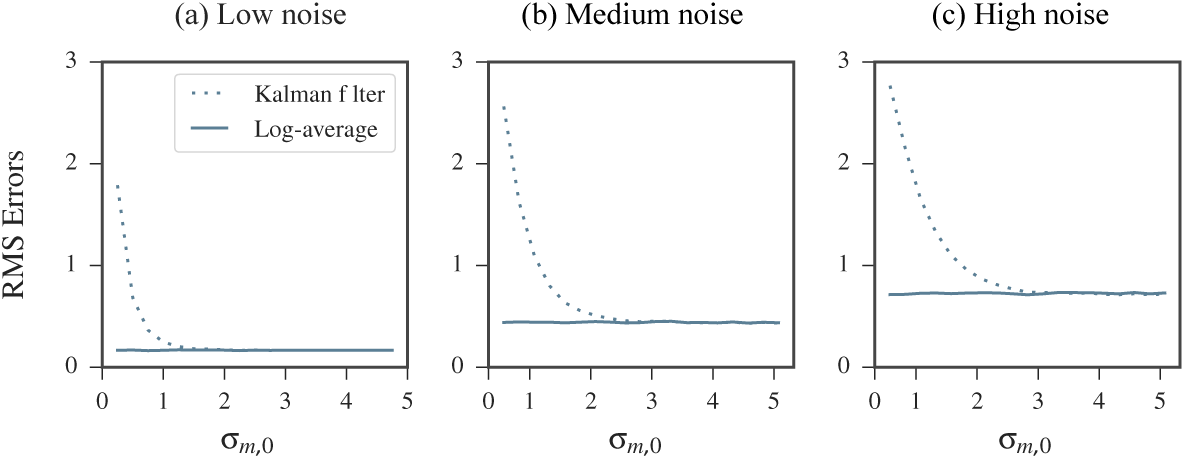
Kalman filtering results using an inaccurate (biased) prior improves with increased uncertainty in prior. RMS errors were calculated over all nucleotides in our database. Error calculations were carried out in the log domain and the ground truth values were the log reactivities. See Methods for RMS calculation details. The prior used in the KF was biased by adding the offset *µ*_offset_= 3 to the ideal prior mean. As the standard deviation of the prior, *σ*_*m,*0_, was increased, the filters performance improved, despite the mean offset. On the other hand, when standard deviation was close to 0, the filter is influenced by a narrow, biased prior and produced poor results.

**Fig 8.**
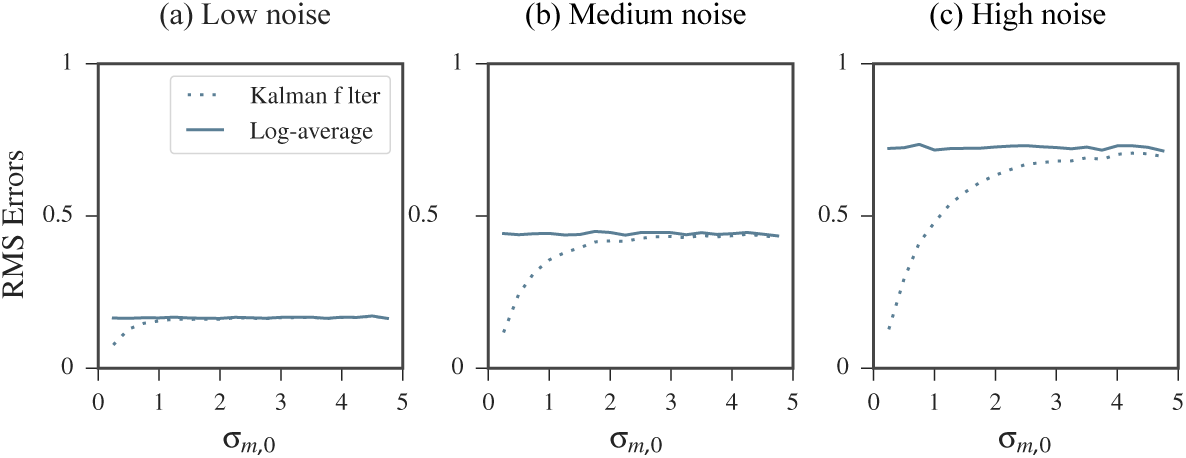
Kalman filtering results using an accurate (unbiased) prior performs comparable to log-averaging when the uncertainty is increased. RMS errors were calculated over all nucleotides in our database. Error calculations were carried out in the log domain and the ground truth values were the log reactivity. See Methods for RMS calculation details. The prior mean was fixed to the ideal value. Its standard deviation, *σ*_*m,*0_, was then increased. As the standard deviation increased, the more comparable the Kalman filtering’s performance was to log-averaging.

As expected, the KF performed best when provided with an accurate and precise prior distribution. Its performance suffered the most when the prior mean offset was increased but its standard deviation remained small. However, when the KF was fed a highly inaccurate but also imprecise prior, the results mirrored that of log-averaging.

While these simulations can be seen as a purely theoretical exercise, we note that the prior distribution was modeled based on data collected from years of RNA SHAPE experiments. As more data is obtained, data characterizations will inevitably improve. It is thus not far-fetched to foresee future datasets that beget more specialized prior models.

### Comparison of data-directed structure predictions under different replicate processing strategies

A major applications of SHAPE data is in RNA secondary structure prediction. In dynamic programming based secondary structure prediction algorithms, reactivities are incorporated into the structure prediction algorithm by first being converted into a pseudo-energy change term. This term is based on a linear-log relationship between reactivities and pseudo-energies. Thus, the prediction algorithm internally transforms the input profile to the log domain. For this section, we employ the RNAstructure software package [55], which implements such an algorithm. When using multiple replicates, the goal is to first combine them in a way that optimally removes the noise component. The resulting profile is then used as input to the prediction software to ultimately improve prediction accuracies. The replicate processing can be done either in the data domain by averaging, or in the log domain by log-averaging or Kalman filtering. To compare these three approaches, we ran the following sets of computational experiments to make secondary structure predictions on each of the 22 RNAs in our database:

1. **Reference set (SET0):** The original SHAPE profile (ground truth) was used as input to RNAstructure. The accuracy of the resulting predicted structure was used as a baseline for comparison to those predicted in SET1, SET2, and SET3.
2. **Average set (SET1):** We generated 3 replicates under the log-normal noise model for each RNA. In the data domain, the average profile was calculated and used as input to RNAstructure.
3. **Log-average set (SET2):** Using the same 3 replicates, the log-average profile was calculated in the log domain, transformed back to the data domain, and used as input to RNAstructure.
4. **Kalman filter set (SET3):** Using the same 3 replicates, the KF profile was calculated in the log domain, transformed back to the data domain, and used as input to RNAstructure.

For each set, the differences between the predicted structure and the reference structure were quantified using the Matthews Correlation Coefficient (MCC) [56, 57] (see Methods for MCC definition). As SET0 is the baseline set, we subtracted the MCC values of SET1, SET2, and SET3 from those in SET0. These results are shown in Fig. 9 for 3 replicates simulated in the low, medium, and high noise regimes. Results using 2 and 4 simulated replicates are shown in S1 Fig and S2 Fig. For replicates simulated under moderate noise levels, we did not observe substantial differences between the results of SET1, SET2, and SET3. However, in the presence of high noise, the structures predicted in SET2 and SET3 (using the log-average and KF profiles, respectively) were closer in MCC to the baseline than SET1 (using the average profile). Comparing the results of SET1 (averaging) and SET2 (log-averaging), for 17 of the 22 RNAs, the MCC coefficients for the structures predicted using the log-average profiles were closer to the baseline than those predicted using the average profiles. For these RNAs, the improvement observed in the results in SET2 compared to SET1 was between 0.69% and 48.21%. For the remaining RNAs, the decrease in MCC values in SET2 compared to SET1 was less than 6.05%. On the other hand, the differences between the results of the two log-domain processed profiles, SET2 (log-averaging) to SET3 (Kalman filtering) where negligible, even in the high noise regime.

**Fig 9.**
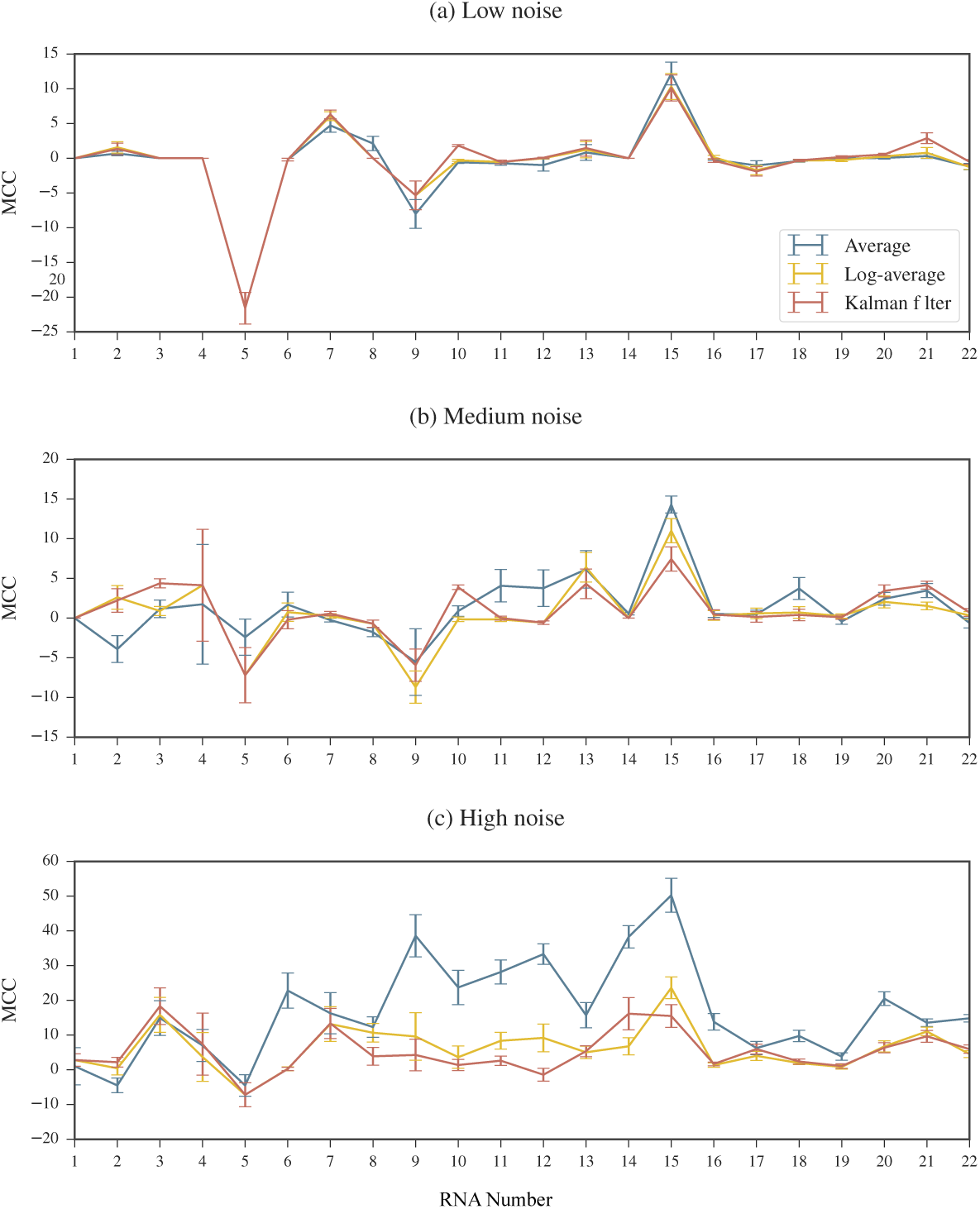
RNAstructure results for profiles calculated using different processing methods. 3 replicates simulated at (a) low (b) medium and (c) high noise regimes. MCC differences are plotted compared to the baseline calculated in SET0. An MCC difference of 0 indicates that when the processed profile was used as input to the RNAstructure software, the resulting predicted structure had the same accuracy as the one predicted using the ground truth profile as input. A positive MCC difference indicate that when the processed profile was input to to the RNAstructure software, the resulting predicted structure was less accurate than the one predicted using the ground truth profile as input. Note that the scale of the MCC differences vary between noise regimes. RNAs are ordered by length. See Table 1 of Methods for corresponding sequence names and lengths. Error bars represents standard errors over 10 repeated runs of replicate simulations.

## Discussion

In this work, we explored models of noise in SHAPE experiments and compared methods for replicate processing. The goal of replicate processing is to generate a profile that captures as well as possible the true sequence of reactivities. This is done by combining measurements for each nucleotide in a way that eliminates the contaminating noise. Any statistically sound processing method is closely linked to the model describing the system. A system model includes models for both the reactivity of a nucleotide and the noise effecting measurements, which is composed of many contributing factors. Based on an empirical distribution of SHAPE data, we modeled reactivities as following a log-normal distribution. We described two models for the measurement noise in SHAPE experiments: the normal noise model and the log-normal noise model. In both models, each nucleotide in an RNA was assumed to have a ground truth reactivity value that persists between replicates. Nucleotide reactivities were also assumed to be independent across an RNA. Considering the normal noise model, replicate processing corresponds to simple measurement averaging. In the log-normal noise model, we outlined two methods for replicate processing: log-averaging and Kalman filtering. Our analyses of SHAPE experiments underscored that a normal noise model is not adequate to represent the data. We instead discussed the relevance of the log-normal noise model. Under the assumptions of this model, we noted that processing such experiments by data domain averaging leads to bias in the resulting profile. This bias can have an affect on the ensuing applications of the data, such as in the case of data-directed RNA secondary structure prediction. These detrimental effects can be avoided by carrying out the replicate processing in the log domain, either by log-averaging or Kalman filtering. Within the log-normal noise model, application of the Kalman filtering approach has the advantage that a prior on the nucleotide reactivities can be introduced. The performance of Kalman filtering is directly dependent upon the quality of the prior and replicate processing can significantly improve with a reliable prior. This auxiliary prior information employed by the filter is particularly useful for signal extraction in the case of substantial noise or as the number of replicates decreases. Accordingly, a well characterized prior represents an additional opportunity for improvement in signal extraction beyond data quality and replicate count.

As mentioned above, Kalman filtering results are strongly tied to the quality of this prior. We observed that a high quality prior mitigates the use of multiple replicates, which can be a serious advantage in resource limited analysis of large RNA molecules. Because such a prior is based on an empirical distribution which can be built with any reasonably sized database, we take this opportunity to advocate the use of public data. As more data becomes available, we anticipate that more specialized priors can be generated, further improving filtering results. We again note that although we focused on the SHAPE probe in this work, there are a variety of other experimental probes available providing a wealth of opportunity for data characterization.

## Future directions

Kalman filtering is just one of many possible signal processing methods available for information extraction. In fact, the KF is a specialized form of the general class of Bayesian filters [58]. Extended Kalman filters and particle filters and other members of this class of filters loosen the Kalman constraints and can also be applied to the analysis of SHAPE data.

A distinct advantage of filtering is that, as with the use of the prior distribution, it provides opportunity to incorporate other types of information into the denoising scheme. Consider, as one example, the correlation effects of neighboring nucleotides in SHAPE experiments, which have been noted and modeled [54]. Although in our study we assumed independence between nucleotides, these effects can be incorporated into processing algorithms to improve signal extraction. Such complex modeling is simply inaccessible under an averaging framework, leaving these correlations as untapped avenues for improved signal extraction.

As a final note, we reiterate that much work is to be done to fully characterize the noise in any SP experiment. The intimate coupling of noise characterization and signal extraction underscores the importance of this step in data processing. Although structure prediction is the most prominent applications of SHAPE data, there exists a breadth of emerging applications for SP data, such as data-directed sequence alignment and the identification of conserved and functional RNA structures [27, 54, 59]. SP data and filtering techniques need to be examined in the context of these data-drive applications.

## Materials and Methods

### Preprocessing SHAPE data

Normalized SHAPE reactivity scores are expected to fall between 0 and 2. However, values exceeding 2 and below 0 are not rare and most SHAPE profiles contain both negative and 0 values. Thus, prior to the application of a log transformation, the profile must undergo some preprocessing. A common approach for dealing with negative values is to simply replace each occurrence with 0 [33]. We refrained from using this method as a profile processed in this way still precludes the use of the log transformation. Another approach is to replace negative reactivities with their absolute value. The drawback of this approach stems from the distribution of negative valued reactivities: while negative values correspond to unreactive nucleotides, the long tail in the distribution can result in an unreactive nucleotide being assigned an uncharacteristically high reactivity.

To circumvent these problems, we followed a procedure similar to the one taken in [30]. Using a large set of SHAPE data, we built a “background distribution” from the empirical distribution of all negative values observed. Our background distribution included data coming from the SHAPE profiles of all 22 RNAs in our database (see Table 1 of Methods). All values below a certain cutoff were removed from this set in order to truncate the tail of the background distribution. In our experiments, we set this cutoff to −0.25. For a given profile, each negative and 0 valued reactivity were replaced by sampling from the truncated distribution. The absolute value of this sample was used as the updated reactivity. After all negative and 0 valued reactivities were replaced, the resulting processed profile was strictly positive and amenable to a log transformation. The original and processed SHAPE profiles of the 22 RNAs in our database are included in S1 Dataset.

### Simulation of replicates

To generate a replicate under the log-normal noise model for an RNA with ground truth profile *S*, we simulated the reactivity measurements for each nucleotide *m* separately. As log measurements follow Eq. 3, the log reactivity of nucleotide *l*_*m*_ is corrupted by additive noise *w*_*m*_ following distribution 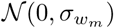. A log measurement was simulated by sampling from this distribution and adding it to *l*_*m*_. We selected 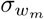 from a uniform distribution *𝒰* (*σ*_min_, *σ*_max_). The values of *σ*_min_ and *σ*_max_ were dictated based on the selected noise regime (See Results for definition of noise regimes). This was repeated for the *M* nucleotides in the RNA sequence to generate a complete replicate profile. Replicates were reverted to the data domain via an exponential transformation. A comparison of the mean-dependence in the standard deviation of real and simulated replicates are shown in Fig. 10.

**Fig 10.**
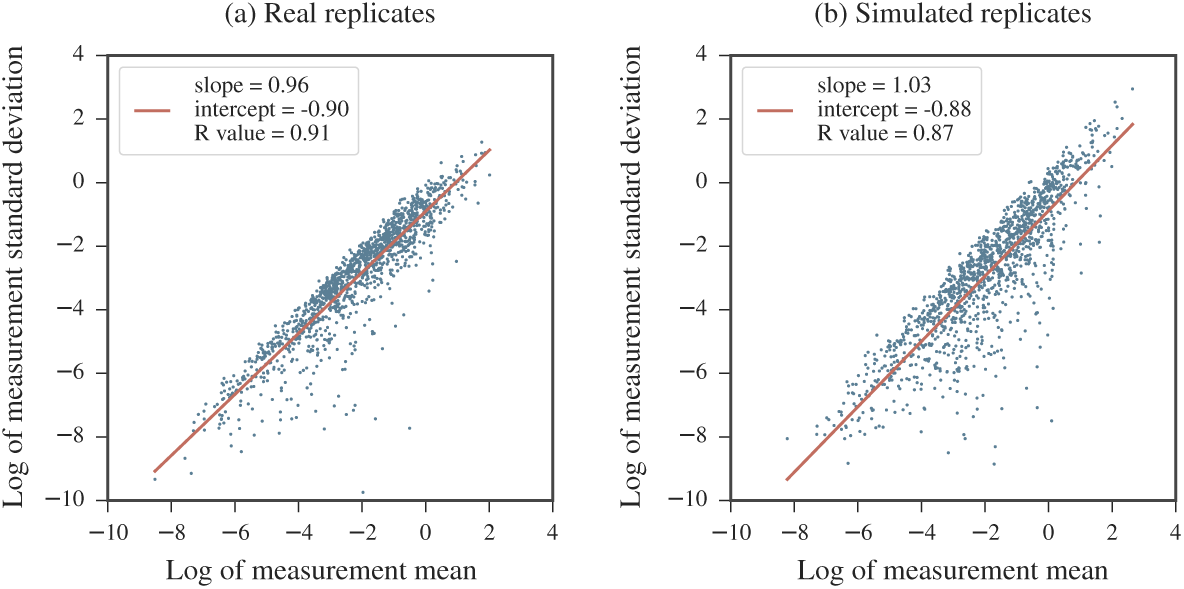
Comparison of mean-dependence in the standard deviation of (a) real and (b) simulated SHAPE measurements. For each nucleotide, the mean value of the 5 measurements (real and simulated) were calculated and plotted against their standard deviation on a log-log plot. A linear fit is overlaid in red for each. The left panel is a recreation of Fig. 1 for comparison. The right panel consists of data coming from simulated replicates for the same RNA. The ground truth reactivity used the in replicate simulation was the average measurement per nucleotide coming from the real replicates. For the simulated replicates, noise levels were between *σ*_min_ = 0 and *σ*_max_ = 1.5. Note that negative reactivity values in the real data are not included as they are incompatible with the log-log plot.

### Kalman filter implementation

We now provide a description of the simplified 1 dimensional implementation of the KF we applied in the log domain. To maintain notational simplicity in this section, we drop the *m* subscripts denoting the nucleotide but restate that the filter is applied per nucleotide.

Recall in the log domain the relationship between the log measurements *l*_*i*_ and the true log reactivity *l* is

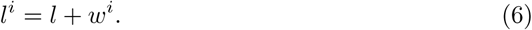

We assume the *w*^*i*^ values are independent and identically distributed as ∼ *𝒩* (0, *σ*_*w*_). The measurement vector is [*l*^1^, *l*^2^,*…, l*^*N*^]. The order of measurements imposed in this vector is random and does not affect the final filtered result. The variance, 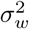, represents the uncertainty in each measurement. Its approximate value, 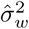, is the sample variance of the *l*^*i*^ values. That is,

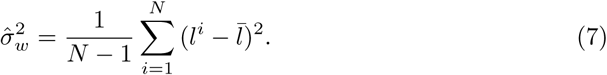

The prior distribution is denoted *𝒩* (*µ*_0_, *σ*_0_). The log reactivity for a nucleotide, *l*, is assumed to be a sample of this distribution. We set *µ*_0_ = *-*1.74 and *σ*_0_ = 1.52. These values were obtained using Gaussian fit to the empirical distribution of our database of 10690 log transformed SHAPE reactivity values. Let 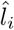 denote the optimal estimate of *l* after the *i*^th^ KF iteration. The uncertainty in this estimate is denoted by 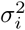. The Kalman gain term at the *i*^th^ iteration is denoted by *K*_*i*_.

The filter is initialized as follows. Prior to the inclusion of the first measurement, the estimate 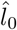 relies solely on the prior. The estimate is thus the prior mean and its uncertainty is the same as the prior variance. That is,

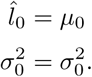

During the *i*^th^ KF iteration, the *i*^th^ measurement, *l*^*i*^, is incorporated into the estimate. First, the Kalman gain is calculated as:

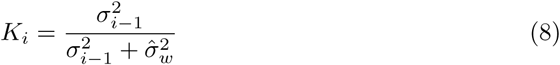

The new estimate, 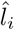, and its uncertainty, *σ*_*i*_, are then calculated as:

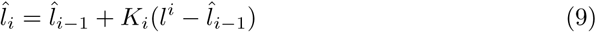

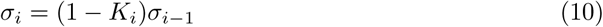

The uncertainty, *σ*_*i*_, is in fact the variance in the posterior distribution of the prior conditioned on the measurements incorporated so far. This value decreases as more measurements are incorporated. The new estimate represents an optimal fusion of the previous estimate and the newly incorporated measurement. The filter repeats Eqs. 8 - 10 until all *N* measurements have been incorporated into the model. The final estimate of *l* is 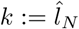.

Note that our implementation appears to bypass the predict step of the standard KF algorithm. This is because we assume no uncertainty in our model that the nucleotide’s reactivity remains constant between replicates. Thus, the predicted value for the (*i* + 1)^st^ measurement is simply the *i*^th^ estimate, 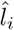.

A Python implementation of this method is provided in S1 File.

### Ideal prior for the Kalman filter

The ideal prior is perfect information. Such a prior has a mean that is the value to be predicted and a standard deviation of 0. For a nucleotide *m* with ground truth reactivity *s*_*m*_, the prior distribution used in the KF is denoted *𝒩* (*µ*_*m,*0_, *σ*_*m,*0_). In the case of the ideal prior, *µ*_*m,*0_ = *l*_*m*_ and *σ*_*m,*0_ = 0. We studied how deviations from this ideal model affected the KF results by adding an offset to the ideal mean. That is,

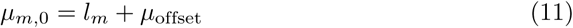

The offset value, *µ*_offset_, was varied between −3 and 3. The prior standard deviation, *σ*_*m,*0_, which signifies the uncertainty in the prior, was similarly increased from 0 to 5.

### Error calculations

We calculated the root mean square (RMS) error over all nucleotides considered (in an RNA or relevant bin for heat map generation) as

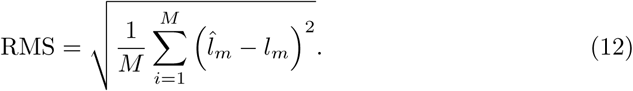

Here, 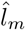 is the value to be compared against the ground truth, *l*_*m*_. *M* is the number of nucleotides considered (in an RNA or relevant bin for heat map generation). For our calculations, 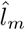 was either the log-average reactivity, 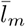, or the KF reactivity, *k*_*m*_.

### Matthews Correlation Coefficient

The accuracy of a computationally predicted secondary structure for a given RNA sequence can be assessed by comparing it to a reference structure. The number of true positives, TP, is the number of base pairs that appear in both structures. The number of false positives, FP, is the number of base pairs that appear in predicted structure but not in the reference structure. The number of true negatives, TN, is the number of possible base pairs that do not appear in either structure. Finally, the number of false negatives, FN, is the number of base pairs that appear in the reference structure but do not appear in the predicted structure. As defined in [57], the MCC value of the predicted structure is calculated as

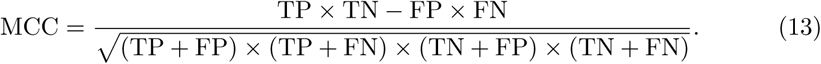

### Data used

Fig. 1, 3, and 10 were created using the cucumber mosaic virus RNA3 sequence data from [47]. The database used in the rest of our analysis was comprised of data coming from the 22 RNAs listed in Table 1 with their appropriate source. The total number of nucleotides in our database was 11070. From the published SHAPE profiles of these RNAs, 1262 of the nucleotides have non-positive SHAPE reactivities. These were used to build the background distribution described above. Another 380 nucleotides do not have SHAPE scores recorded in the published profiles. Hence, a total of 10690 SHAPE reactivities were used in our study.

**Table 1.**
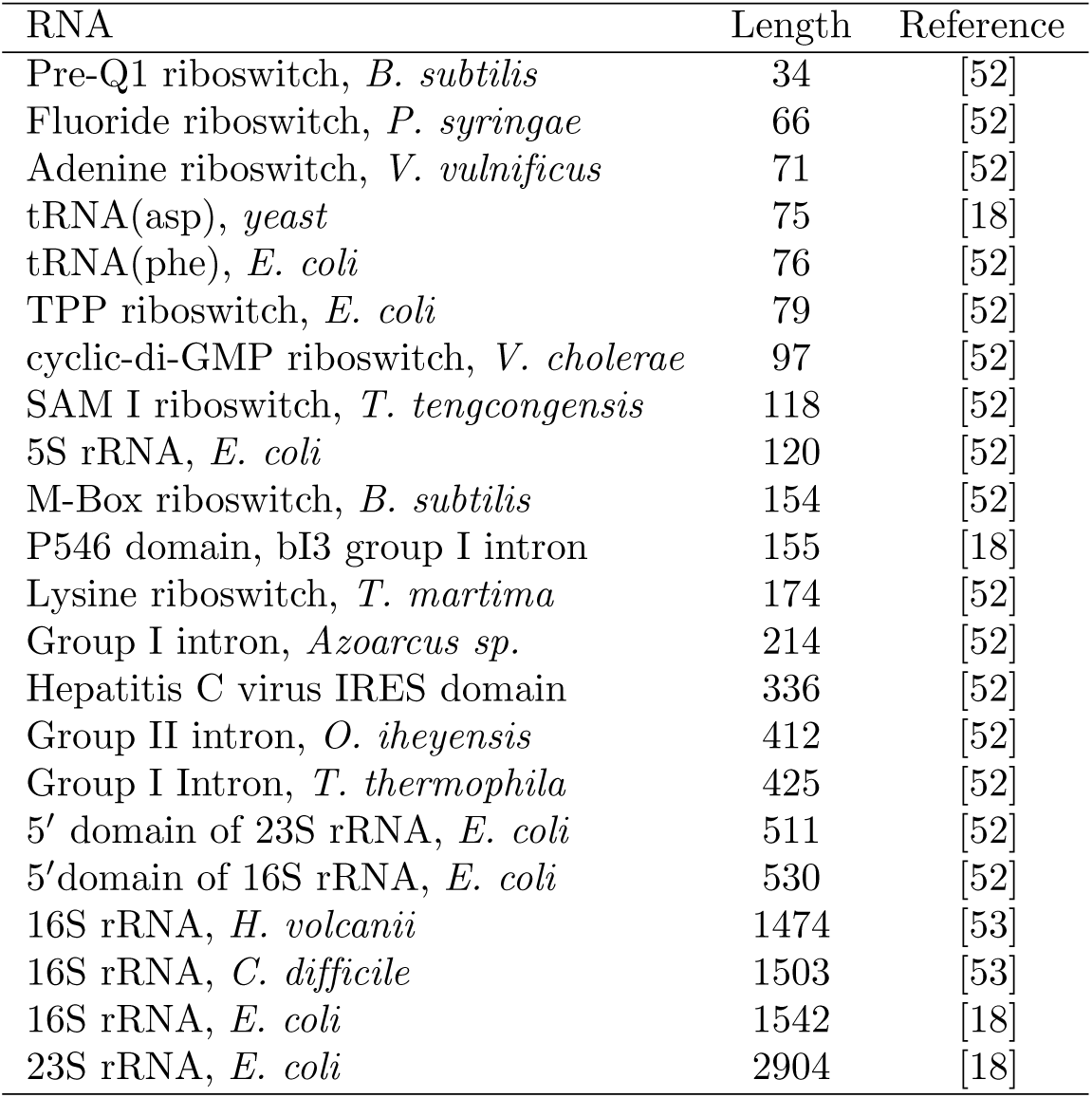
Summary of RNA sequences with SHAPE profiles included in database.

## Supporting information

**S1 Fig. RNAstructure results for profiles calculated using different processing methods.** 2 replicates simulated at (a) low (b) medium and (c) high noise regimes. MCC differences are plotted compared to the baseline calculated in SET0. An MCC difference of 0 indicates that when the processed profile was used as input to the RNAstructure software, the resulting predicted structure had the same accuracy as the one predicted using the ground truth profile as input. A positive MCC difference indicate that when the processed profile was input to to the RNAstructure software, the resulting predicted structure was less accurate than the one predicted using the ground truth profile as input. Note that the scale of the MCC differences vary between low and high noise regimes. RNAs are ordered by length. See Table 1 of Methods for corresponding sequence names and lengths. Error bars represents standard errors over 10 repeated runs of replicate simulations.

**S2 Fig. RNAstructure results for profiles calculated using different processing methods.** 4 replicates simulated at (a) low (b) medium and (c) high noise regimes. MCC differences are plotted compared to the baseline calculated in SET0. An MCC difference of 0 indicates that when the processed profile was used as input to the RNAstructure software, the resulting predicted structure had the same accuracy as the one predicted using the ground truth profile as input. A positive MCC difference indicate that when the processed profile was input to to the RNAstructure software, the resulting predicted structure was less accurate than the one predicted using the ground truth profile as input. Note that the scale of the MCC differences vary between low and high noise regimes. RNAs are ordered by length. See Table 1 of Methods for corresponding sequence names and lengths. Error bars represents standard errors over 10 repeated runs of replicate simulations.

**S1 Dataset. Original and processed SHAPE profiles for the 22 RNAs of Table 1.**

**S1 File. Python implementation of 1D Kalman filter for RNA SHAPE replicates.**

## Acknowledgments

This work was supported by National Institutes of Health (NIH) grants: HG006860 from the National Human Genome Research Institute (NHGRI)-NIH and T32-GM008799 from The National Institute of General Medical Sciences (NIGMS)-NIH. Its contents are solely the responsibility of the authors and do not necessarily represent the official views of the NIGMS or NIH.

